# Successful exome capture and sequencing in lemurs using human baits

**DOI:** 10.1101/490839

**Authors:** Timothy H. Webster, Elaine E. Guevara, Richard R. Lawler, Brenda J. Bradley

**Affiliations:** School of Life Sciences, Arizona State University, Tempe, AZ 85287; Center for the Advanced Study of Human Paleobiology, The George Washington University, Washington, DC 20052; Department of Anthropology, Yale University, New Haven, CT 06511; Department of Sociology and Anthropology, James Madison University, Harrisonburg, VA 22807

**Keywords:** genomics, strepsirrhines, primates, macaques, methods

## Abstract

**Objectives:** We assessed the efficacy of exome capture in lemurs using commercially available human baits.

**Materials and Methods:** We used two human kits (Nimblegen SeqCap EZ Exome Probes v2.0; IDT xGen Exome Research Panel v1.0) to capture and sequence the exomes of wild Verreaux’s sifakas (*Propithecus verreauxi,* n = 8), a lemur species distantly related to humans. For comparison, we also captured exomes of a primate species more closely related to humans (*Macaca mulatta,* n= 4). We mapped reads to both the human reference assembly and the most closely related reference for each species before calling variants. We used measures of mapping quality and read coverage to compare capture success.

**Results:** We observed high and comparable mapping qualities for both species when mapped to their respective nearest-relative reference genomes. When investigating breadth of coverage, we found greater capture success in macaques than sifakas using both nearest-relative and human assemblies. Exome capture in sifakas was still highly successful with more than 90% of annotated coding sequence in the sifaka reference genome captured, and 80% sequenced to a depth greater than 7x using Nimblegen baits. However, this success depended on probe design: the use of IDT probes resulted in substantially less callable sequence at low-to-moderate depths.

**Discussion:** Overall, we demonstrate successful exome capture in lemurs using human baits, though success differed between kits tested. These results indicate that exome capture is an effective and economical genomic method of broad utility to evolutionary primatologists working across the entire primate order.

## Introduction

Recent advances in next generation sequencing technology and the increasing availability of annotated reference genomes have made feasible the genomic study of nonmodel taxa (Ellegren, 2014; Goodwin, McPherson, & McCombie, 2016). Nonhuman catarrhines, in particular papionin monkeys (Bergey, Phillips-Conroy, Disotell, & Jolly, 2016; Gibbs et al., 2007; Lea, Altmann, Alberts, & Tung, 2016; Wall et al., 2016) and apes (Carbone et al., 2014; de Manuel et al., 2016; Locke et al., 2011; Perry et al., 2008; Prado-Martinez et al., 2013), have been the focus of intense genomic study because of their importance in understanding human evolutionary history (Jolly, 2001; Swedell & Plummer, 2012; Wrangham, 1987) and history of use as biomedical models (Carlsson, Schapiro, Farah, & Hau, 2004; Rogers & Gibbs, 2014; Varki, 2000). However, genomic data hold promise to enable vast insights into evolution, ecology, and behavior, as well as inform conservation management across the entire primate order.

Nevertheless, genomic analyses remain out-of-reach for many species. Even for species for which there is a draft genome available, population-scale whole genome sequencing and the concomitant data storage, management, and analyses often require prohibitively vast financial, computational, and bioinformatics resources. These conditions have fostered the development and wide adoption of reduced representation genomic sequencing methods, like restriction-associated DNA sequencing (RAD-seq; K. R. Andrews, Good, Miller, Luikart, & Hohenlohe, 2016; Baird et al., 2008). While RAD-seq and similar “genotyping-by-sequencing” methods have enabled the genomic study of a variety of nonmodel organisms, aspects of the data—particularly marker sparseness and discontinuity—can be limiting for some research questions (Arnold, Corbett-Detig, Hartl, & Bomblies, 2013; Lowry et al., 2017; Rubin, Ree, & Moreau, 2012).

In contrast, targeted capture involves the selective enrichment of genomic regions before sequencing, allowing both for more continuous sequence and for control over the density and identity of targets (Gnirke et al., 2009; Jones & Good, 2016). Foremost among targeted capture techniques is exome capture and sequencing (exome sequencing), which primarily targets all the protein coding regions of the genome along with a number of untranslated regions, promoter regions, and miRNAs (Clark et al., 2011). In total, these targets account for less than 2% of the genome, making exome sequencing much more cost-effective than whole genome sequencing, while still providing the majority of data often desired by those undertaking high throughput sequencing. Numerous commercial exome capture kits based on the human genome have been developed and widely adopted in clinical settings and for identifying the underlying basis of human genetic disorders (Bamshad et al., 2011; Bilgüvar et al., 2010; Ng et al., 2010).

Synthesizing custom high-quality oligonucleotide baits for targeted capture is expensive and generally requires a high-quality reference genome (Jones & Good, 2016; but see Snyder-Mackler et al., 2016). Because of the close evolutionary, and thus genetic, relationship between human and nonhuman primates, researchers studying nonhuman primates are advantageously situated to potentially exploit the baits and resources developed for human exome sequencing. In particular, human exome baits have been successfully used in haplorrhine primates (Bataillon et al., 2015; George et al., 2011; Hvilsom et al., 2012; Jin et al., 2012; Teixeira et al., 2015; Vallender, 2011). However, it is currently unclear how well human exome baits would work for more distantly related species (e.g., strepsirrhine primates).

To ascertain and quantify the utility of exome sequencing across the order Primates, we performed exome capture and sequencing of a distantly related strepsirrhine species, Verreaux’s sifaka (*Propithecus verreauxi*), that diverged from humans over 60 million years ago (dos Reis et al., 2018). As a direct comparison to provide context for assessing the strepsirrhine results we also included rhesus macaques (*Macaca mulatta*), a catarrhine species for which the efficacy of exome capture using baits designed for humans has already been established (George et al., 2011; Vallender, 2011). Both species have closely-related reference genomes available (*P. coquereli, M. mulatta*). Our overall goal is to assess capture efficiency, mapping success, and variant calling using two commercially available human exome capture kits.

## MATERIALS AND METHODS

### Samples

We collected the Verreaux’s sifaka samples from individuals living at Bezà Mahafaly Special Reserve (Bezà), located in southwestern Madagascar (Toliara province). As part of long-term research, research team members capture unmarked yearlings and recent immigrants annually to collect biometric data and give each individual a unique identifying collar and ear notch pattern (Richard, Dewar, Schwartz, & Ratsirarson, 2002).

For this study, we generated two different Verreaux’s sifaka datasets (Sifaka1 and Sifaka2). Sifaka1 is the primary dataset we use throughout the study in comparison with the macaque samples (Macaque1). We generated the Sifaka2 dataset using different exome capture kit to explore any effects of bait design on capture success (described below). For Sifaka1, we extracted DNA from banked ear tissue biopsies as described in Lawler et al. (2001) from four sifakas: a mother-daughter pair and two unrelated males (Supporting Information Table S1). For Sifaka2, we extracted DNA from the ear tissue of two additional male and two additional female sifakas using the QIAgen DNeasy Blood and Tissue (Qiagen) kit following manufacturer instructions with an extended lysis step (Supporting Information Table S1).

For a catarrhine comparison, we used DNA derived from blood samples from four unrelated—two male and two female—captive Indian rhesus macaques (Macaque1) from the Wisconsin National Primate Research Center (Supporting Information Table S1).

### DNA extraction, library preparation, and sequencing

We sent extracted DNA to the Yale Center for Genome Analysis (YCGA) for exome capture, library preparation, and multiplexed sequencing following their standard protocols, described as follows. For all three datasets (Sifaka1, Sifaka2, and Macaque1), genomic DNA was sheared to a mean fragment length of 140 bp and adapters were ligated onto both ends of fragments. Fragments were then PCR amplified, during which a 6 bp barcode was inserted at one end of each fragment. Libraries were hybridized with baits from two different kits: Nimblegen baits (Nimblegen SeqCap EZ Exome version 2) were used for Sifaka1 and Macaque1, and IDT xGen baits (IDT xGen Exome Research Panel 1.0) were used for Sifaka2. Fragments were then mixed with streptavidin-coated beads and washed to remove unbound fragments. Captured fragments were then PCR amplified and purified with AMPure XP beads. Libraries from Sifaka1 and Macaque1 were multiplexed (all four sifaka samples in one lane, and the four macaques in another lane with two other samples) and sequenced using 75 bp paired-end reads on a single lane of an Illumina HiSeq 2000 using Illumina protocols. Sifaka2 libraries were sequenced using 100 bp paired-end reads on a single Illumina HiSeq 4000 lane and multiplexed with eight other samples (12 total samples per lane, but only four are included in this study).

### Exome assembly

We assessed read quality pre- and post-trimming using FastQC (S. Andrews, 2018) and MultiQC (Ewels, Magnusson, Lundin, & Käller, 2016). We used BBDuk (Bushnell, 2018) to remove adapters and perform quality trimming using the parameters “ktrim=r k=21 mink=11 hdist=2 tbo tpe qtrim=rl trimq=10”. We then mapped reads from sifaka samples (Sifaka1 and Sifaka2) to the *Propithecus coquereli* draft genome (Pcoq_1.0; Baylor College of Medicine; https://www.ncbi.nlm.nih.gov/genome/24390). P. verreauxi and P. coquereli share a common ancestor 3-8 million years ago (Herrera & Dávalos, 2016; Springer et al., 2012). We mapped macaque samples (Macaque1) to the Indian rhesus macaque draft genome (Mmul_8.0.1; Macaca mulatta Genome Sequencing Consortium; https://www.ncbi.nlm.nih.gov/genome/215?genome_assembly_id=259055). Finally, we mapped reads from both species to the human reference genome (hg38; Genome Reference Consortium, Dec 2013). For the rest of the manuscript, we refer to Mmul_8.0.1 as mmul8, proCoq_1.0 as pcoq1, and hg38 as hg38. In all cases, we mapped reads using BWA MEM (Li, 2013) using default parameters except for “-t 4” and “-R” to add read group information. We marked duplicates with SAMBLASTER (Faust & Hall, 2014). We then used SAMtools (Li et al., 2009) to fix read pairing, and sort and index BAM files.

To enable a direct comparison of exome capture success between species for which we had different numbers of raw reads and different duplication rates, we conducted all downstream analyses on downsampled BAM files (containing the same number of reads for each individual). To downsample BAM files, we first used the “stats” tool in SAMtools (Li et al., 2009) to count the total number of reads and number of duplicate reads in each BAM file. We then used the “view” tool in SAMtools (Li et al., 2009) with the parameters “-F 1024 –s 0.<subsample_fraction>” to subsample approximately 50 million reads, where <subsample_fraction> is equal to 50 million divided by the total number of nonduplicate reads. The flag “-F 1024” removes reads flagged as duplicate.

### Variant calling

We jointly called variants for each dataset using both GATK’s HaplotypeCaller (Poplin et al., 2018) and Freebayes (Garrison & Marth, 2012). To speed up processing, we input BED files containing minimally callable sites—depth greater than 3, mapping quality greater than 19, and base quality greater than 29—generated using CallableLoci in GATK (McKenna et al., 2010). Finally, we filtered variants for site quality (minimum of 30), sample depth (minimum of 8), sample genotype quality (minimum of 30), allele support (minimum of 3 reads), and number of passing samples (minimum of 4) with a Python script built using the cyvcf2 library (Pedersen & Quinlan, 2017).

We functionally annotated filtered variants using Ensembl’s Variant Effect Predictor (McLaren et al., 2016) tool with annotations derived from the NCBI gene format files corresponding to the respective reference genomes for the rhesus and sifaka references (NCBI *Macaca mulatta* Annotation Release 102 [GCF_000772875.2] and *Propithecus coquereli* Annotation Release 100 [GCA_000956105.1]), and Ensembl’s cache for the human reference (hg38). We also obtained NCBI’s annotation for the human reference (GCF_00001405.37) for use in our coverage analyses (see below). Using these annotation files, we intersected various regions (exon, intron, and intergenic) with filtered variants using bedtools “intersect” (Quinlan & Hall, 2010) and then used the “stats” module of BCFtools (Li, 2011) to tally variants in each region.

### Coverage analysis

We calculated the mean and standard deviation of mapping quality (MAPQ) of reads within each BAM file using a custom program written in Go (*“mapqs.go”*) using packages in bíogo/hts (Kortschak, Pedersen, & Adelson, 2017). BWA MEM’s (Li, 2013) MAPQ scores are PHRED-scaled and can range from 0-60, with higher values indicating increased confidence in mapping accuracy.

We counted the number of callable sites across a variety of depths and genomic regions by first using SAMtools (Li et al., 2009) “view” to remove duplicates and reads with a mapping quality less than 20 with the flags “-F 1024 -q 20”, and then calculating per site depths with *genomecov* in bedtools (Quinlan & Hall, 2010), outputting in bedgraph format (“-bg”). We then processed bed files, including intersecting with genomic regions derived from the NCBI annotation described above using bedtools (Quinlan & Hall, 2010), BEDOPS (Neph et al., 2012), and a custom Python script (“*Compute_histogram_from_bed.py”*). Finally, we used the *coverage* module in bedtools with default parameters (Quinlan & Hall, 2010) to calculate the fraction of each coding region with coverage. Note that for all region-based analyses, we merged regions in the NCBI GFF annotations during processing because many, but not all, regions were present multiple times.

### Exome capture kit comparison

We used the sifaka datasets (Sifaka1 and Sifaka2) for a direct comparison of capture success using the two different capture kits (NimbleGen SeqCap EZ Exome version 2 for Sifaka1 and IDT xGen Exome Research Panel 1.0 for Sifaka2). We ran both datasets through identical exome assembly and coverage analysis steps as described above.

### Data Availability

We deposited raw sequencing reads in NCBI’s Sequence Read Archive (https://www.ncbi.nlm.nih.gov/sra) under BioProject PRJNA417716. We provide SRA accession numbers in Supporting Information Table S1.

We built all analyses into a reproducible pipeline using Snakemake (Köster & Rahmann, 2012), Bioconda (Grüning et al., 2018). The entire pipeline—including all scripts, environment files, and software versions—is available on Github (https://github.com/thw17/Sifaka_assembly).

### Ethics Statement

We report no conflict of interest. All research conformed to institutional and national guidelines, and complied with the American Association of Physical Anthropologists Code of Ethics. This protocol is approved by the James Madison University Institutional Animal Care and Use Committee (protocol numbers A03-14 and A18-04) and permission to conduct research at Bezà was granted by the Malagasy Ministry of the Environment.

## 3. Results

We generated the following mean numbers of raw sequencing reads per sample: 96,358,883 for Sifaka1 (range 80,434,390–108,538,766), 65,460,924 for Macaque1 (range 59,422,700–70,596,638), and 75,441,014 for Sifaka2 (range 70,264,610– 79,520,110) (Supporting Information Table S2). After trimming, duplicate removal, and quality control, 83-86% of Sifaka1 reads, 89-92% of Macaque1 reads, and 68-70% of Sifaka2 reads passed all filters, with differences among datasets largely driven by duplication rates (Supporting Information Table S2). To account for these differences in duplication rates and raw sequences generated, we downsampled reads for all samples to approximately 50 million nonduplicate reads (Supporting Information Table S2). We only included the downsampled datasets in downstream analyses.

Mapping qualities were very similar when mapping samples to their most closely related reference genomes (pcoq1 for sifakas; mmul8 for macaques). Across datasets, we observed mean mapping qualities of approximately 56 (out of a maximum of 60), with standard deviations ranging between 11 and 14 (Figure 1). However, when mapping to the human reference genome (hg38), mapping qualities decreased substantially—dropping to approximately 52 in Macaque1, 45 in Sifaka1, and 48 in Sifaka2—and the standard deviation increased (Figure 1).

**Figure 1.**
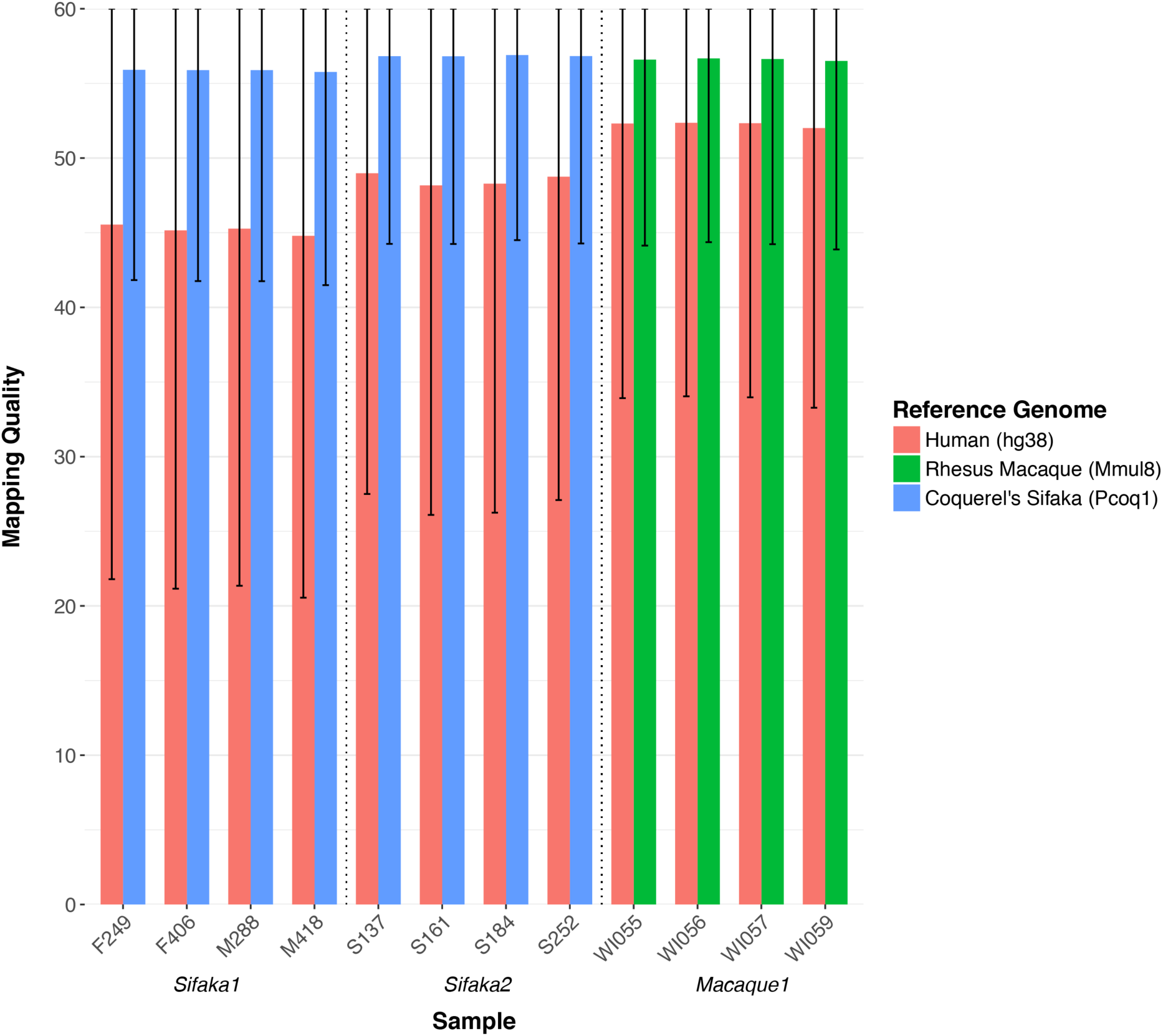
Mean mapping quality (MAPQ) for samples mapped to their most closely related reference genome (pcoq1 for sifakas and mmul8 for macaques) and the human reference genome (hg38). Samples are organized by dataset membership, defined by species and capture kit. Sifaka1 and Macaque1 were processed using NimbleGen baits and Sifaka2 was processed using IDT baits. Error bars denote plus/minus one standard deviation. Note that maximum mapping quality is 60.

We measured the number of sites in coding (CDS), intergenic, intronic, and untranslated (UTR) regions at four different depth thresholds (1x, 4x, 8x, and 12x), counting only nonduplicate reads with a minimum mapping quality of 20, which we term “callable sites.” Across all regions and in both datasets (Sifaka1 and Macaque1), we observed a decrease in the number of callable sites as we increased minimum depth of coverage (Figure 2). This decrease was minor for CDS and UTR, while intronic and intergenic regions exhibited a disproportionate drop moving from 1x to 4x thresholds (Figure 2). We observed taxon differences as well. Specifically, the Macaque1 samples exhibited more callable sites in each region than those in Sifaka1 for all reference genomes. Moreover, we found little difference between callable sites in mmul8 and hg38 for each region in Macaque1, in contrast to Sifaka1, for which we observed a decrease in callable sites across regions when moving from pcoq1 to hg38 (Figure 2).

**Figure 2.**
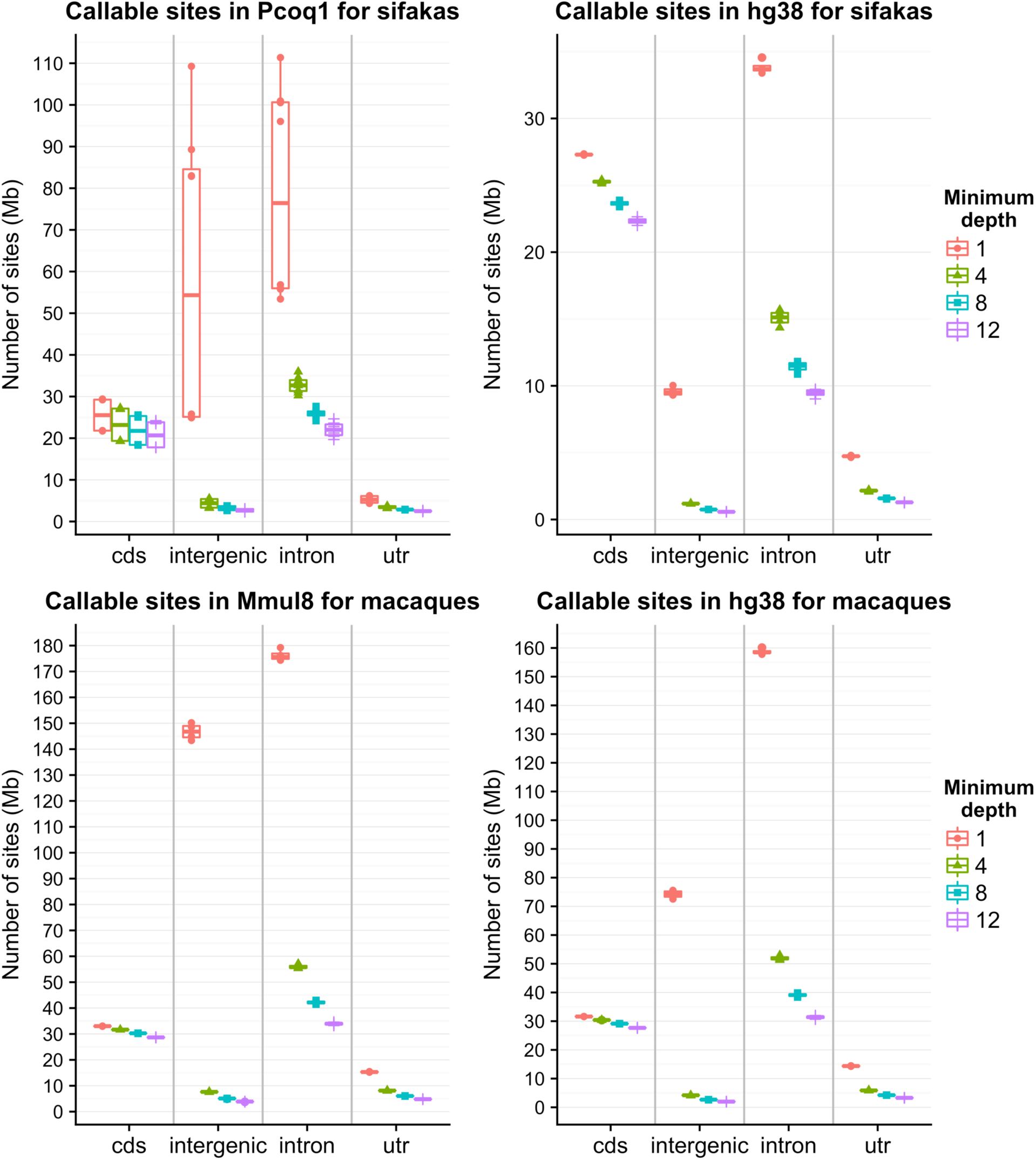
The effect of minimum depth on the number of callable sites across genomic regions. Samples are mapped to their most closely related reference genome (pcoq1 for sifakas and mmul8 for macaques) and the human reference genome (hg38). Minimum depth thresholds were 1, 4, 8, and 12 nonduplicate reads per site with MAPQ greater than or equal to 20.

Because the primary goal of exome sequencing is to target coding sequence, we explored CDS in more detail (Figure 3; Figure 4). For both Sifaka1 and Macaque 1, when mapping to the most closely related reference genome (pcoq1 and mmul8, respectively), we found that more than 90% of annotated CDS had one or more reads mapped to it (Figure 3; Sifaka1 mean = 90.9%, Macaque1 mean = 92.8%). However, as the minimum depth threshold increased, we observed a steeper decline in Sifaka1 than Macaque 1 until approximately 20x coverage. For example, Sifaka1 had means of 84.1% (4x), 78.7% (8x), 74.0% (12x), 69.9% (16x) and 66.1% (20x) of CDS covered at increasing thresholds, while Macaque1 had broader coverage at each threshold: 89.1% (4x), 85.1% (8x), 80.7% (12x), 75.8% (16x), and 70.6% (20x) of CDS covered (Figure 3). This pattern was far more pronounced when the two datasets were mapped to hg38. Across the same depth thresholds, Sifaka1 had approximately 10-14% fewer bases covered when mapping to pcoq1 to hg38 (Figure 3), while Macaque1 only exhibited a 3- 5% decrease per threshold moving from mmul8 to hg38 (Figure 3).

**Figure 3.**
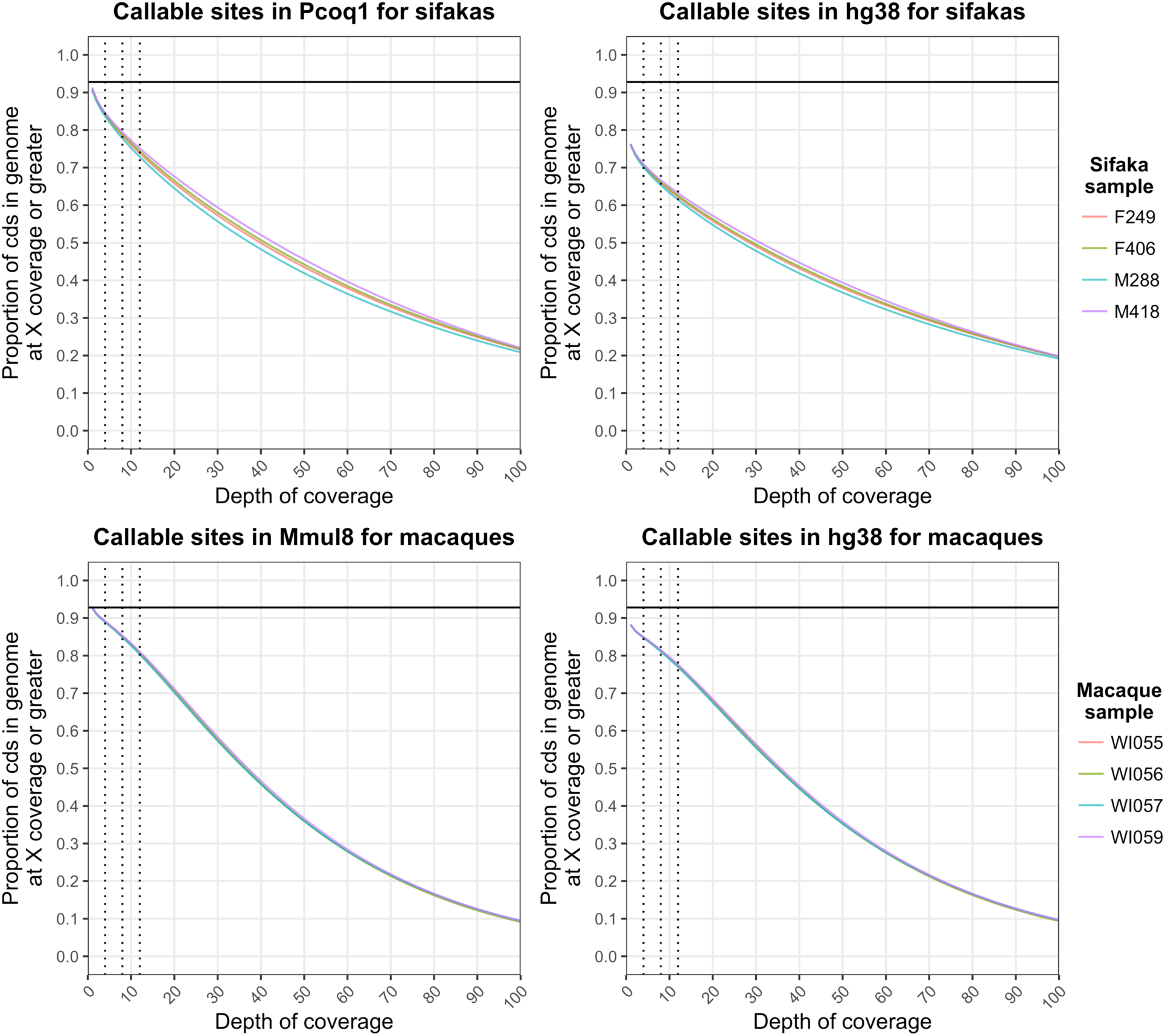
Depth of coverage across the coding regions of the genome. Samples are mapped to their most closely related reference genome (pcoq1 for sifakas and mmul8 for macaques) and the human reference genome (hg38). The x-axis presents depth of coverage, measured as the number of nonduplicate reads with MAPQ >= 20. The y-axis presents the proportion of coding sequence in the genome with X or greater coverage, where X is the value on the x-axis. The vertical dotted lines highlight three common filter values: 4x or greater coverage, 8x or greater coverage, and 12x or greater coverage. The solid horizontal line marks the fraction of the genome covered by one or more reads for macaque samples mapped to mmul8.

**Figure 4.**
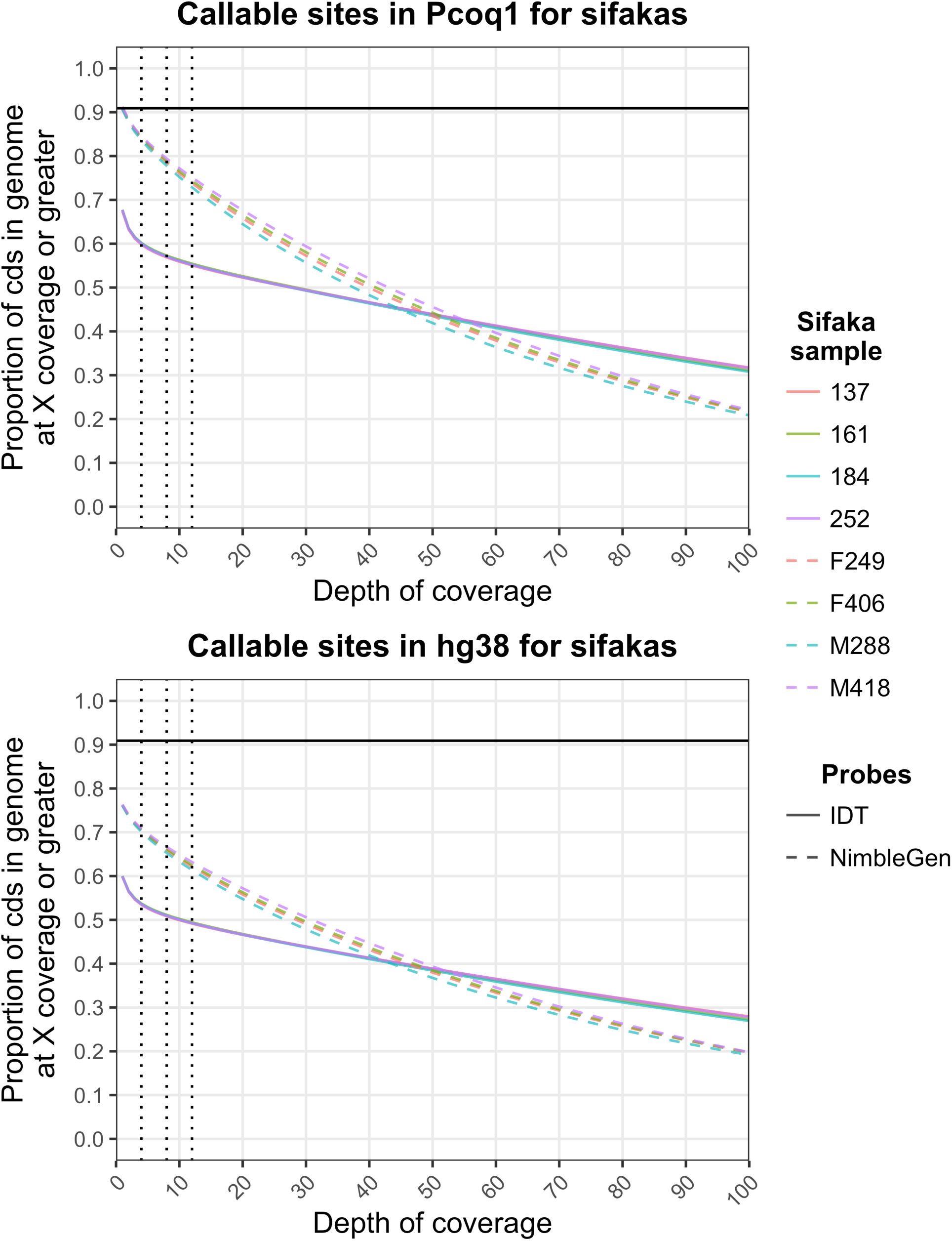
A comparison of depth of coverage across the coding regions of the genome for NimbleGen and IDT baits. Samples are mapped to pcoq1 and hg38. Samples captured with NimbleGen baits are represented by dashed lines, while those captured with IDT baits are represented by solid lines. The x-axis presents depth of coverage, measured as the number of nonduplicate reads with MAPQ >= 20. The y-axis presents the proportion of coding sequence in the genome with X or greater coverage, where X is the value on the x-axis. The vertical dotted lines highlight three common filter values: 4x or greater coverage, 8x or greater coverage, and 12x or greater coverage. The solid horizontal line marks the fraction of the genome covered by one or more reads for NimbleGen samples mapped to pcoq1.

We also tested to see if exome capture success in strepsirrhines was consistent across two commonly used commercially available human kits: NimbleGen (Sifaka1) and IDT (Sifaka2). At lower minimum depth thresholds typically used in genomic analyses (e.g., 8x and 12x), the NimbleGen kit recovered more than 20% more CDS in pcoq1 and 15% more CDS in hg38 than IDT (Figure 4). This difference was significant across depths less than 50x (U= 31378, p < 2.2 x 10^−16^). Interestingly, because NimbleGen and IDT exhibit different slopes, they intersect at approximately 50x coverage (Figure 4). While NimbleGen probes still recover significantly more CDS at depth thresholds between 50x and 100x (U=38416, p < 2.2 x 10^−16^), the proportion of bases with *X* or more coverage exhibits the opposite pattern in this interval, with IDT displaying higher values (Figure 4). This pattern is consistent with IDT capturing less sequence, but at greater depths (i.e., depths greater than 100x).

To further explore the difference in capture success between NimbleGen and IDT, we calculated the breadth of coverage across coding regions in pcoq1. NimbleGen probes captured a significantly greater mean fraction of coding regions, measured as the mean fraction of each coding sequence covered by at least one read (185,162 regions; NimbleGen mean = 0.91, IDT mean = 0.63; U=1.89 x 10^11^, p < 2.2 x 10^−16^). Upon closer examination, this difference was primarily driven by IDT completely missing more coding regions. Among 185,162 coding regions in pcoq1, 33.8% of regions lacked coverage in the IDT data (range 33.6-34%), while only 7.1% completely lacked coverage in the NimbleGen dataset (range = 6.6-7.3%). When we excluded these regions with no coverage, the difference in mean fraction of coding regions captured decreased substantially, though NimbleGen still captured significantly more (NimbleGen mean = 0.98, IDT mean = 0.95; U=1.57 x 10^11^, p < 2.2 x 10^−16^).

We used two variant callers, GATK’s HaplotypeCaller and Freebayes, to genotype Sifaka1 and Macaque1 when mapped to hg38 and the closest reference (pcoq1 for Sifaka1, and mmul8 for Macaque1), for a total of eight sets of variant calls (Supporting Information Table 3). In both datasets, variant call sets for the most closely related genome were broadly similar between HaplotypeCaller and Freebayes in terms of number of variants identified and genes overlapped (Supporting Information Table 3). However, when mapped to hg38, the datasets showed opposite patterns: for Sifaka1, HaplotypeCaller identified approximately four times as many variants as Freebayes (HaplotypeCaller = 250,389, Freebayes = 62,793), while in Macaque1 Freebayes identified more than 56% more variants (HaplotypeCaller = 97,709, Freebayes = 152,768; Supporting Information Table 3).

While the number of variants identified across call sets differed substantially, within call sets, proportions of variant types were broadly similar (Supporting Information Table 3). Most variants identified across call sets were single nucleotide variants (SNVs; 71.6-81.2%), though proportions of multiple nucleotide variants (MNVs), insertions, and deletions increased when mapping to hg38. Similarly, the relative numbers of nonsynonymous, frameshift, and stop gained variants in exons were much higher when mapping to hg38 (Table 1). Variants were not limited to exons however, as most variants were intronic (Supporting Information Table 3).

**Table 1.** Coding variants identified.^a^

| Coding variant type | Sifaka1<br>(*Pcoq1*) | Sifaka1<br>(*hg38*) | Macaque1<br>(*Mmul8*) | Macaque1<br>(*hg38*) |
| --- | --- | --- | --- | --- |
| Synonymous | 62,201 | 114,951 | 96,650 | 95,982 |
| Nonsynonymous | 29,079 | 102,711 | 51,743 | 75,499 |
| Frameshift variant | 1,230 | 9,648 | 1,044 | 13,171 |
| Stop | 409 | 3,563 | 408 | 3,231 |

<sup>a</sup>Values are counts of variants identified for each dataset-reference genome combination.

## 4. Discussion

In this study we demonstrate, for the first time, that human baits can be used to successfully capture high-coverage exomic data for strepsirrhines. While previous studies have established that human baits are effective in anthropoid primates (George et al., 2011; Jin et al., 2012; Vallender, 2011), our results extend the cross-species application of baits to lineages diverged over 60 million years ago (dos Reis et al., 2018) and indicate that human baits are likely viable options for genomic analyses across the entire primate order.

We found that a mean of 90.9% of annotated coding sequence (CDS) in the draft *P. coquereli* genome was covered by one or more reads in our *P. verreauxi* samples. As we increased the minimum depth of coverage thresholds to match common filter values (e.g., 4x, 8x, and 12x coverage), we observed a steady, curvilinear decline in breadth of CDS coverage (Figure 3). This pattern indicates that coverage is not uniform across CDS, consistent with predictions for next-generation sequencing (Lander & Waterman, 1988), particularly those for targeted capture (Clark et al., 2011; Sims, Sudbery, Ilott, Heger, & Ponting, 2014). In particular, in targeted sequencing, there are expected position-based sampling biases that lead to greater coverage towards the middle of targets (Wendl & Barbazuk, 2005). However, despite the fact that increasing sequencing effort will increase depth nonuniformly across targets, clearly any CDS base with coverage has been successfully captured. Therefore, increasing sequencing effort—we used 50 million nonduplicate reads in this study—should increase the fraction of callable CDS at various coverage thresholds up to at least 90.9%, the amount of CDS we observed covered by at least one read in this study.

Surprisingly, the fraction of captured CDS in sifakas (90.9%) was very similar to, albeit slightly smaller than, the fraction captured in rhesus macaques (92.8%), even though macaques share a much more recent common ancestor with humans (30-35 million years; dos Reis et al., 2018). However, the macaques exhibited a slower decrease in breadth of CDS coverage at increasing minimum depth thresholds, particularly across thresholds most commonly used (Figure 3). Thus, while exome capture is certainly highly successful in sifakas, there is a decrease in efficiency compared to lineages more closely related to humans. This pattern holds both when mapping to the nearest reference genomes (pcoq1 for the sifakas and mmul8 for the macaques) and when mapping back to the human reference, and therefore appears to be driven by capture success, rather than assembly methods (e.g, mapping).

Similar to our results, previous studies have noted a decrease in capture efficiency across increasing evolutionary distances within catarrhine primates (Jin et al., 2012; Vallender, 2011). In this study, however, we found that this effect was much less pronounced even though we sampled much greater evolutionary distances. This is likely driven by differences in mapping strategies. Previously, assessments of capture efficiency involved mapping back to the human reference genome (George et al., 2011; Jin et al., 2012; Vallender, 2011). In this study, we found that mapping across large evolutionary distances appears to reduce both breadth and depth of coverage (Figure 3), an effect likely caused by the greater number of differences between reads and the reference sequence, which substantially impacts mapping quality (Figure 1). In fact, the Indian rhesus macaques used in this study were much more closely related to their nearest reference (same population and species) than the Verreaux’s sifakas (about 6 million years diverged from *P. coquereli*; dos Reis et al., 2018), which might account for some of the difference in our observed capture success between the two species, though this requires further study. In addition, while most protein-coding genes in sifakas and macaques are expected to have homologues in humans, gene content is not identical across primates (Rogers & Gibbs, 2014). It is therefore possible that our results were also influenced by the presence of more sifaka-specific gene content than macaque-specific gene content.

While exome capture was successful in the sifakas, the degree of success depended on the capture baits used. Specifically, while the NimbleGen probes captured an average of more than 90% of pcoq1 CDS and only completely missed 6-7% of coding regions, the IDT probes captured less than 70% of pcoq1 CDS and completely missed approximately one-third of coding regions. When we excluded missed regions, the difference in coverage reduced substantially, with both baits covering more than 95% of CDS in regions with any coverage. Taken together, the difference between baits is primarily driven by IDT baits completely missing entire coding regions, rather than the failure of IDT baits to capture entire targets. Commercially available human exome capture kits differ markedly in design, with different targets, bait lengths, and bait overlap (Clark et al., 2011). Even in human samples, for which the baits are designed, these differences in bait design affect capture efficiency and the number and location of variants detected (Clark et al., 2011; Sulonen et al., 2011).

Compared to other reduced representation methods (e.g., RAD-seq), exome capture’s primary strength is that it aims to capture all protein coding regions of the genome—the regions frequently of most interest from a functional standpoint. To this end, exome capture and sequencing, particularly with the NimbleGen probes, was highly successful in our samples, capturing the vast majority of CDS and leading to the identification of a rich suite of variants. However, exome capture’s utility is not limited to these regions, and it can generate high-quality data in regulatory and untranslated regions (UTRs), as well as other intronic and intergenic regions (Samuels et al., 2013). In our data, we identified tens of millions of base pairs of sequence outside of coding regions (Figure 2); in fact, more variants were identified in introns than any other sequence class. Thus, exome capture across nonhuman primates holds great promise for not only recovering coding regions across the genome, but also recovering putatively neutral sequences (introns, intergenic regions, and four-fold degenerate sites) that can be applied to traditional questions in molecular ecology regarding kinship, geneflow and demographic history.

## Acknowledgments

We thank Sibien Mahereza, Enafa Jaonarisoa, Elahavelo Efitroarane, Efitiria, Edouard Ramahatratra, Alison Richard, Roshna Wunderlich, Jeannin Ranaivonasy, and Joel Ratsirarson for helping with sample collection at Bezà Mahafaly; Roger Wiseman (Wisconsin National Primate Research Center) for providing macaque samples; and Gary Aronsen for lab assistance. We are grateful to the Nacey Maggioncalda Foundation (THW); National Science Foundation (NSF BCS-1455818; THW); the Yale Institute for Biospheric Studies (THW), Yale University (BJB); The George Washington University (EEG, BJB); the Yale MacMillan Center (EEG), and the Wenner-Gren Foundation (EEG) for generous financial support. Many analyses were conducted on the Louise and Ruddle High Performance Computing Clusters at Yale University (supported by NIH grants RR19895 and RR029676-01) and the Arizona State University High Performance Computing Cluster.

