## Supporting Information Table 1 for "Successful exome capture and sequencing in lemurs using human baits"

| Sample | Species | Origin | Dataset | Sex | SRA Accession |
| --- | --- | --- | --- | --- | --- |
| F249 | *P. verreauxi* | Bezà | Sifaka1 | Female | SRR6277165 |
| F406 | *P. verreauxi* | Bezà | Sifaka1 | Female | SRR6277166 |
| M288 | *P. verreauxi* | Bezà | Sifaka1 | Male | SRR6277167 |
| M418 | *P. verreauxi* | Bezà | Sifaka1 | Male | SRR6277168 |
| S137 | *P. verreauxi* | Bezà | Sifaka2 | Female | SRR7219233 |
| S161 | *P. verreauxi* | Bezà | Sifaka2 | Male | SRR7219232 |
| S184 | *P. verreauxi* | Bezà | Sifaka2 | Male | SRR7219231 |
| S252 | *P. verreauxi* | Bezà | Sifaka2 | Female | SRR7219230 |
| WI055 | *M. mulatta* | WiNPRC | Macaque1 | Male | SRR5085658 |
| WI056 | *M. mulatta* | WiNPRC | Macaque1 | Female | SRR5085657 |
| WI057 | *M. mulatta* | WiNPRC | Macaque1 | Female | SRR5085688 |
| WI059 | *M. mulatta* | WiNPRC | Macaque1 | Male | SRR5085648 |
